# MRI-Compatible Rigid Head Holders for Artifact-Free Multimodal Imaging in Mice

**DOI:** 10.64898/2026.05.29.728741

**Authors:** Severin Filser, Daniel Peter Varga, Gian Marco Calandra, Peter Makra, Felix Christopher Nebeling, Bernd Evers-Dietze, Welf Wawers, Arthur Liesz, Paolo Salomoni, David Hecker, Hardik Doshi, Eugenio Fava, Cornelia Hesse, Nikolaus Plesnila, Hans Fried

**Author notes:** These authors contributed equally to this study (Severin Filser, Daniel Peter Varga, Gian Marco Calandra).

## Abstract

**Purpose:** High-resolution intravital microscopy allows cellular-scale analysis of the brain *in vivo* but is greatly sensitive to physiological motion. Combining optical microscopy with magnetic resonance imaging (MRI) in the same animal could relate cellular and mesoscale functional readout to whole-brain structural information, but this requires head holders that are both mechanically rigid and MRI compatible. Conventional metallic head holders introduce MRI artifacts, whereas many nonmetallic alternatives lack sufficient stability for chronic microscopy. Thus, we developed rigid, MRI-compatible head holders engineered from 3D-printed zirconia ceramics to reduce motion during microscopy while preserving MRI image quality.

**Methods:** Head holders were designed for mouse cranial fixation and fabricated from zirconia ceramics using additive manufacturing. We quantified motion artifacts during two-photon and multimodal widefield imaging of the mouse cortex and assessed their impact on neuronal calcium activity, functional connectivity, and hemodynamic readouts. MRI compatibility was evaluated by measuring image quality in the presence of the head holder.

**Results:** The ceramic head holders provided mechanical stability to reduce motion artifacts to micrometer levels during intravital imaging. The head holders produced no detectable susceptibility artifacts in MRI, and image contrast was comparable to control acquisitions performed without head holder. Sequential optical and MRI imaging of the same brain regions established artifact-minimized multimodal data acquisition within the same animal.

**Conclusions:** Non-metallic ceramic head holders support longitudinal multimodal studies that combine high-resolution optical microscopy with whole-brain MRI measurements in the same animal.

## Introduction

*In vivo* light microscopy has revolutionized preclinical neuroscience by providing high-resolution access to cellular, synaptic, and circuit-level processes in the living mammalian brain. Two-photon microscopy, in particular, is a powerful intravital tool for examining neuronal morphology, dendritic spine dynamics, and neuronal activity at cellular resolution in the cortex, with imaging depths that can exceed 400 micrometers [1]. However, their high magnification makes these techniques highly susceptible to motion artifacts resulting from respiration, cardiac pulsation, and spontaneous locomotor activity [2, 3]. Even in anesthetized animals, breathing-induced shifts can significantly degrade image quality and compromise morphological and quantitative analyses [4-6]. These limitations have driven extensive technological development in strategies to suppress motion artifacts, including rigid head fixation, organ immobilization, and image stabilization techniques [7-9].

While intravital microscopy provides exceptional spatiotemporal resolution within a comparatively small accessible volume, its utility in preclinical neuroscience is limited by restricted translatability to clinical settings. Magnetic resonance imaging (MRI), by contrast, has become the gold standard for non-invasive structural and functional assessment of the whole brain across species, providing three-dimensional anatomical detail, quantitative perfusion measurements with arterial spin labeling, diffusion-weighted imaging of white and grey matter microstructure, and blood-oxygen-level-dependent (BOLD) functional mapping [10]. Sequential or simultaneous multimodal imaging combining high-resolution optical techniques with MRI would allow researchers to connect mechanistic insights from microscopy to clinical investigation, while simultaneously reducing animal use through integrated longitudinal studies in individual subjects [11].

However, the combination of intravital microscopy and MRI in the same animal presents substantial technical challenges. Intravital microscopy requires absolute mechanical stability to eliminate motion artifacts during image acquisition, necessitating rigid fixation of the animal’s head through a mounted head holder or restraint apparatus. Simultaneously, MRI imposes stringent compatibility requirements: ferromagnetic and conductive materials proximal to the imaged tissue generate severe susceptibility artifacts through localized magnetic field distortions, which can lead to large brain regions becoming uninterpretable [12, 13]. These metal-induced artifacts scale with magnetic field strength and are particularly problematic in small-animal imaging systems operating at high field strengths (7–11.7 Tesla), where spatial resolution demands make signal loss and distortions especially detrimental [14-17].

Here, we present MRI-compatible rigid head holders fabricated from 3D-printed zirconia for multimodal preclinical brain imaging. We characterize the MRI properties of the head holder materials, quantify motion artifacts during imaging, and validate MRI image quality in the presence of the device. We demonstrate that properly engineered non-metallic head holders prevent MRI artifacts while providing the mechanical rigidity required for stable high-resolution microscopy. This technical advance removes a major barrier to multimodal preclinical imaging by allowing synaptic, cellular, mesoscale, and whole-brain readouts to be acquired from, and linked within the same animal.

## Results

3D printable polymers, like polyester ester keton (PEEK), have been previously used to design magnetic field proof components for ultra-high field MRI [18]. However, PEEK features at least one order of magnitude lower mechanical rigidity compared to metallic or ceramics components (Table S1). Therefore, we set out to evaluate 3D-printable metals and ceramics for potential imaging artifacts caused by magnetic susceptibility. Materials were scanned at an ultra-high field strength of 11.7 Tesla and three-dimensional datasets were acquired using T1-weighted isotropic Fast Low-Angle Shot (FLASH) pulse sequences. Image analysis revealed that ceramics, similar to PEEK, caused almost no imaging artifacts, while metals induced strong field distortions (Fig. 1A). Among all tested materials, aluminum-toughened zirconium dioxide (AtZ), pure zirconium dioxide (zirconia), and PEEK showed the best performance. Furthermore, µCT analysis of the zirconia sample demonstrated the fidelity of stereolithography based ceramic printing, which was comparable to conventional milling methods (Fig. 1B).

**Figure 1.**
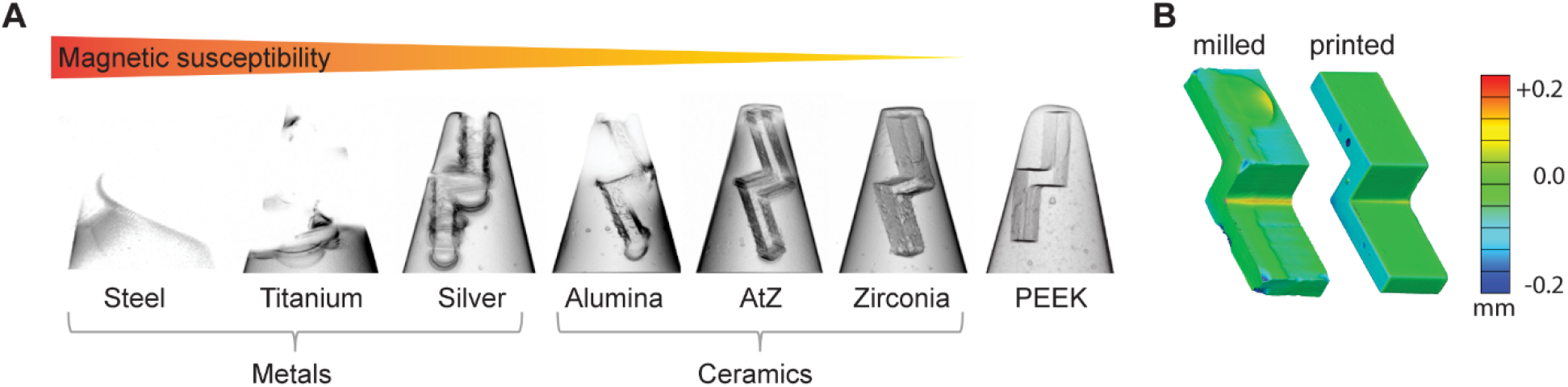
Ceramics produce less MRI artifacts compared to metals at ultra-high field strength (11.7 T). **(A)** µCT surface deviation maps of milled compared to 3D printed zirconia. **(B)** FLASH recordings of agarose embedded materials. Images are arranged from high to low MRI imaging artifacts based on decreasing magnetic susceptibility.

Subsequently, we designed and 3D-printed zirconia head holders (Fig. S1) for mice to immobilize the head during *in vivo* two-photon calcium imaging. We compared the stability of our recordings obtained with ceramic (zirconia) head holders and commercially available standard steel head holders of similar form factor. Awake, head-fixed mice were positioned on a rotating disc treadmill (Gramophone; Femtonics, Budapest, Hungary) to monitor locomotor activity, while neuronal calcium signals were recorded simultaneously from the somatosensory cortex during a non-stimulated baseline period and repeated air-puff stimulation of the whisker pads (Fig. 2A). Air puffs consistently elicited detectable neuronal responses (Fig. 2B, Fig. S2 and movie S1) and significantly increased locomotor activity, as measured by the gramophone (Fig. 2C), providing a suitable testing scenario to evaluate the stabilization performance of the head holder. Image shifts occurred during movement and resting phases (Fig. 2C and D). However, in both scenarios image shifts did not exceed 4 pixels (corresponding to 3 µm) neither in x- or y-direction with a mean displacement of 0.9±1.05 (x-shift) and 0±0.86 (y-shift) for zirconia, and 1.1±2.85 (x-shift) and 1.8±2.01 (y-shift) for steel (Fig. 2D and Fig. S3A). Importantly, average image displacement differed less than 1 µm between steel and zirconia head holders, indicating that zirconia provides sufficient mechanical stability for *in vivo* imaging, in awake, active animals (Fig. 2E).

**Figure 2.**
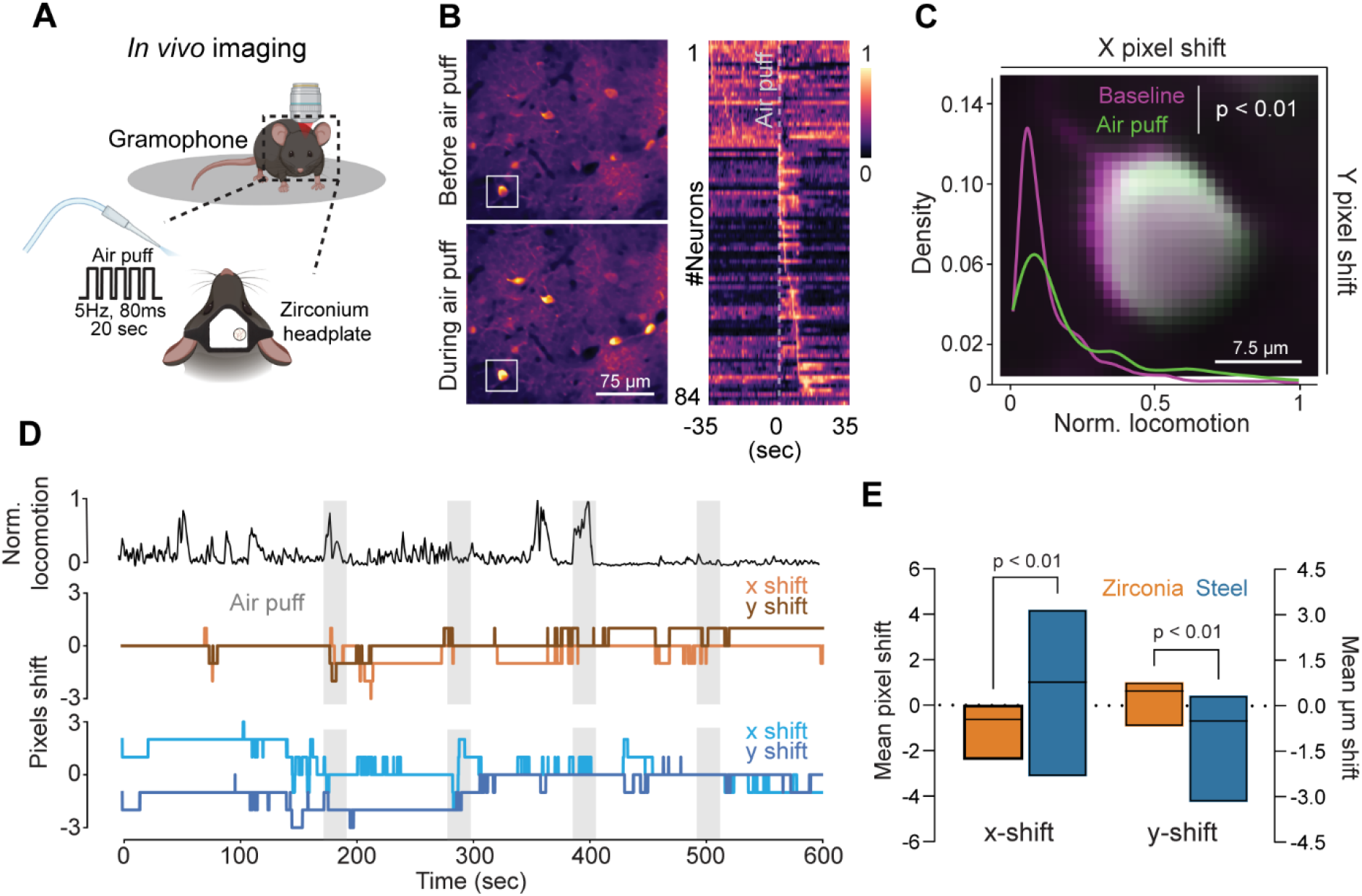
Zirconia head holders enable stable *in vivo* 2-photon recordings of cortical calcium dynamics in awake, mobile mice. **(A)** Schematic of the experimental setup: Experimental animal is head-fixed under a 2-photon microscope using a zirconia head holder and placed on a gramophone to track locomotion. During recording the whiskerpad is stimulated via air puffs for 20 seconds at a frequency of 5 Hz lasting 80 ms per pulse. **(B)** Left: 2-photon micrographs showing cortical calcium responses before (top) and after (bottom) whisker stimulation. Right: Stimulus-aligned activity map showing color-coded calcium responses of detected neurons (n = 84) during the stimulation protocol. **(C)** Density distribution of locomotor activity during baseline (magenta) and stimulation periods (green), together with a representative neuron (enlarged image taken from figure **1B**) illustrating the stimulation-associated shift between the two periods. Density distribution of the data was visualized using a kernel density estimate (KDE), illustrating the probability distribution of values across conditions. Locomotion data from each frame of all zirconia recordings were pooled for comparison (Baseline: 11944 frames; Stimulation: 7968 frames; Mann–Whitney U test). **(D)** Top: Representative locomotion trace recorded during an *in vivo* imaging session, highlighting changes in locomotor behavior during stimulation epochs (grey). Bottom: Representative x/y pixel shift traces from animals implanted with either steel (blue lines) or zirconia (orange lines) headplates. **(E)** Quantification of mean x/y pixel shift during stimulation trials. Data are presented both as pixel shift and corresponding displacement in micrometers. Boxes represent median and extend to the maximum and minimum of the data distribution (Mann-Whitney unpaired *t*-test; Steel vs Zirconia: n = 15 Steel vs 20 Zirconia data points).

Lastly, we used the newly designed zirconia head holders (Fig. S1) to quantify the motion artifacts in awake, head-fixed mice placed on a rotating platform (Gramophone, - Femtonics, Budapest, Hungary) during multimodal mesoscale imaging that combined widefield fluorescence calcium imaging with intrinsic optical imaging (IOI) of hemodynamics. Motion analysis revealed a slightly larger shift along the x axis, perpendicular to the direction of locomotion (Fig. S3, and Movie S2). However, displacement in both axes did not exceed 1 μm. We then evaluated whether residual motion affected subtle region-specific readouts of neuronal activity, functional connectivity, and cerebral blood perfusion under intact and pathological conditions (Fig 3). To relate these functional readouts to whole-brain structural information, we acquired 3T MRI and mesoscale imaging data before and after photothrombotic stroke induction in the left somatosensory cortex (Fig. 3B and F). At 1-day poststroke, lesion formation was confirmed by the reduction of spontaneous calcium activity and IOI signals within the illuminated area, together with T2-weighted MRI evidence of edema extending from the cortical surface to the *corpus callosum* (Fig. 3G, Movie S3). The zirconia head holder was not visible on MRI scans, confirming that it did not introduce detectable imaging artifacts (Fig. 3B and F, Movie S3). Corresponding IOI and calcium recordings revealed spontaneous activity across multiple cortical regions, with region-specific fluctuations occurring on a sub-second timescale (Fig. 3A, Movie S2). Peak calcium responses reached ΔF/F amplitudes of 3 to 4 percentage points. Seed-to-Seed functional connectivity analysis showed robust interhemispheric synchronization between corresponding cortical areas (Fig. 3D, Movie S2). After stroke induction, neuronal calcium activity was markedly reduced in the lesioned cortex relative to the contralateral hemisphere, particularly within the atlas-defined cortical region centered on the photothrombotic induction site (Fig. 3E). Consistently, functional connectivity was strongly disrupted after stroke, with seeds in the ipsilateral hemisphere falling below the correlation threshold (r<0.3) and therefore not displayed in the connectivity map (Fig. 3H). Co-registration of the MRI dataset with IOI-derived hemodynamic map showed close spatial overlap between the hypoperfused cortical area and the MRI-defined lesion (Fig. 3F and G). In parallel, calcium activity in the ipsilateral hemisphere was also suppressed after stroke, although the affected calcium signals reached beyond the corresponding IOI perfusion deficit (Fig. 3G and H). Sagittal and coronal MRI sections revealed substantial three-dimensional edema extending beyond the cortical induction site and involving deeper structures, including white matter and *caudoputamen* (Fig. S4 and Movie S3). These cross-modality comparisons show that widefield calcium imaging and IOI capture cortical functional and hemodynamic consequences of stroke, whereas MRI defines the full three-dimensional extent of tissue involvement. Together, these findings demonstrate that zirconia head holders support chronic, artifact-minimized multimodal *in vivo* imaging and permit direct correlation of mesoscale functional readouts with whole-brain MRI.

**Figure 3.**
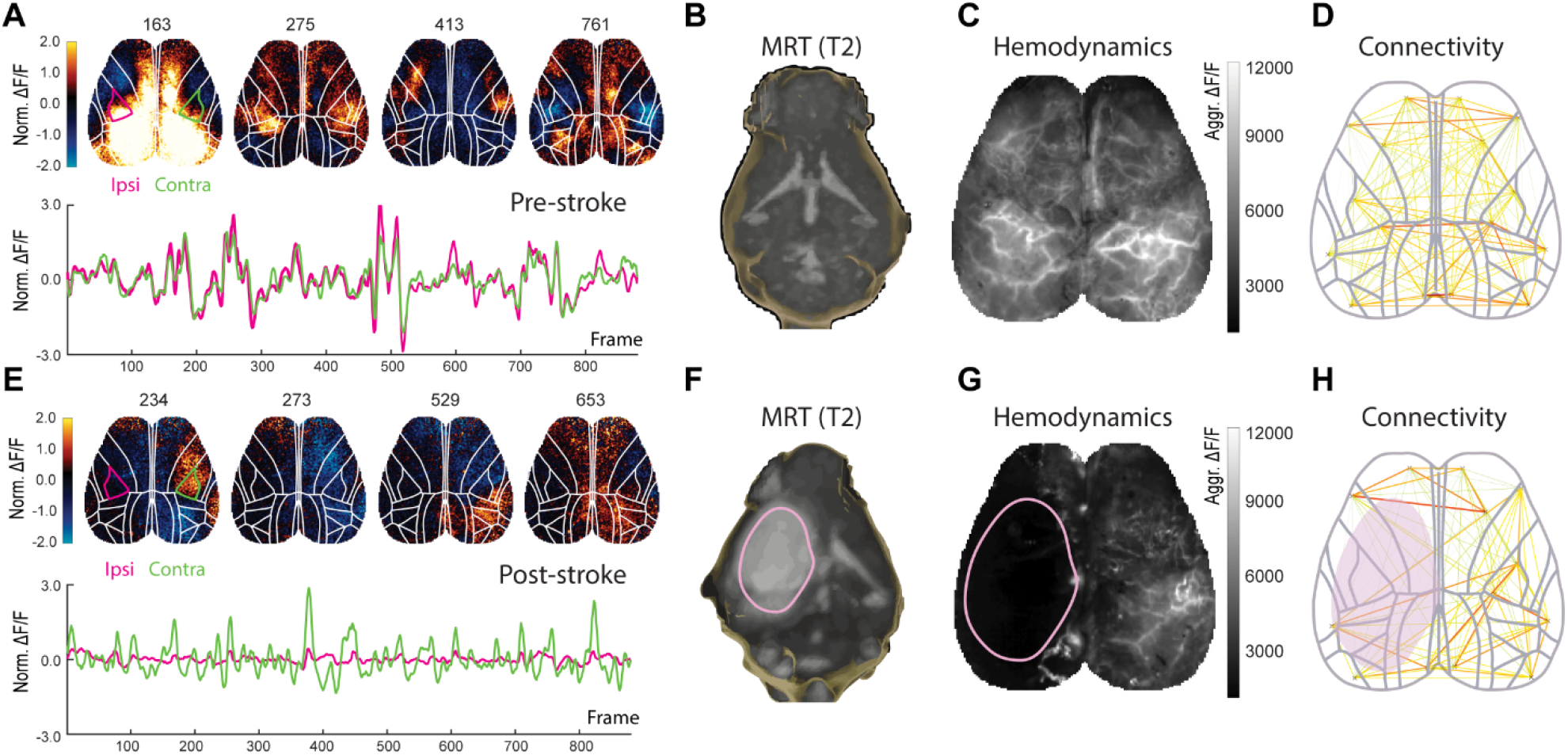
Correlative T2-weighted MRI and multimodal widefield recordings in awake, head-restrained mice before and after photothrombotic stroke induction. **(A and E)** Mesoscale widefield calcium activity before stroke induction **(A)** and after photothrombotic stroke **(E).** Upper subpanels show color-coded, preprocessed widefield calcium signals expressed as ΔF/F percentage-point changes overlaid with cortical boundaries derived from the Allen Mouse Brain Common Coordinate Framework (CCFv3). The ipsilesional region, highlighted in red, corresponds to the center of photothrombotic stroke induction, whereas the homotopic contralesion region is highlighted in green. Lower subpanels show regional calcium time series extracted from these regions under baseline **(A)** and poststroke conditions **(E). (B and F)** T2-weighted MRI images before stroke induction **(B)** and after photothrombotic stroke **(F)**. Images were cropped using an Imaris-derived brain surface mask, shown as a hollow blue outline. In the poststroke image **(F)**, the lesion appears as a hypointense region in the left hemisphere and is delineated by a pink outline derived from Imaris-based three-dimensional segmentation. **(C and G)** Hemodynamic maps derived from green-channel intrinsic optical imaging (IOI) before stroke induction **(C)** and after photothrombotic stroke **(G)**. Maps show aggregated cerebral blood volume– related intensity fluctuations displayed in grayscale. Under baseline conditions **(C)**, cortical vascular architecture is visible, with pial arteries appearing bright and venous structures appearing dark. After stroke induction **(G)**, the ipsilesional cortex shows reduced hemodynamic signal within the edematous tissue, overlapping with the MRI-derived lesion outline shown in pink. **(D and H)** Seed-to-seed functional connectivity maps before stroke induction **(D)** and after photothrombotic stroke **(H)** based on 16 predefined cortical seeds. Lines indicate pairwise connectivity between seeds; thicker red lines denote higher Fisher z-transformed correlation values, whereas thinner yellow-to-green lines denote weaker correlations. Maps are overlaid with atlas-derived cortical boundaries. In the poststroke condition **(H)**, some ipsilesional seeds fell below the correlation threshold and are therefore not displayed. The red area indicates the lesion region defined from the hemodynamic map and MRI-derived segmentation shown in pink.

## Discussion

The development of MRI-compatible rigid head holders manufactured from 3D-printed non-metallic ceramics overcomes a key limitation in correlative preclinical imaging by providing the mechanical stability required for intravital microscopy while maintaining compatibility with of MRI.

Additive manufacturing allows layer-by-layer fabrication of complex geometries directly from CAD models, enabling rapid prototyping and iterative refinement on timescales of days rather than weeks [19-21]. Printing head holders from non-metallic ceramics matches mechanical rigidity with MRI compatibility, as these materials avoid the susceptibility artifacts produced by metals and show magnetic properties similar to biological tissue [22-24]. Furthermore, tested ceramics exhibit stiffness (elastic modulus) values that are at least 50 times higher than those of thermoplastic polymers such as PEEK (Table S1). In addition, zirconia shows approximately twice the fracture toughness of bulk PEEK and up to five times the fracture toughness of 3D-printed PEEK (10 – 13 vs 6.17 and 4.25 MPa *m^1/2^, respectively) [25].

Combining multiple light microscopy approaches with MRI in the same animal over days, and potentially weeks, allows brain structure and function to be examined across spatial and temporal scales that are not accessible with either modality alone. At the micro-scale, two-photon microscopy provides an *in vivo* readout of neuronal structure and physiology, linking structural features such as dendritic spine density with calcium-dependent activity [9]. At the mesoscale level, widefield calcium imaging captures fast cortical dynamics and functional connectivity through genetically encoded calcium indicators, while IOI adds complementary hemodynamic readouts related to cortical blood perfusion [18, 26]. At the whole-brain macro-scale, MRI offers three-dimensional structural information and quantitative measures beyond the optical field of view. This combination is difficult to achieve with conventional steel head holders, which are incompatible with MRI, or with common polymer-based alternatives, which may deform, fracture, or lose mechanical stability during chronic use. In the photothrombotic stroke model, this multiscale approach allowed local cortical changes to be interpreted in the context of whole-brain tissue involvement within the same animal. Notably MRI revealed edema extending far beyond the cortical induction site and involving deeper structures that were not accessible to dorsal optical imaging alone. Repeated acquisition in the same animal facilitates spatial alignment between optical and MRI datasets and allows longitudinal changes to be followed directly.

## Conclusion

Together, these findings show that combining suitable materials with additive manufacturing can overcome competing technical requirements in biomedical imaging. MRI-compatible ceramic head holders produced by 3D printing may enable correlative imaging workflows that link cellular- and mesoscale optical readouts to macroscale whole-brain MRI measures in the same animal *in vivo*.

## Material and Methods

### 3D Printing

Head holders have been 3D printed via lithography-based metal manufacturing (steel - Metshape, Germany), selective metal sintering (titanium, silver - CADdent Prototyping, Germany) and ceramic stereolithography (alumina, AtZ, zirconia - CADdent Prototyping, Germany). PEEK head holders have been produced via mechanical drilling (CADdent Prototyping, Germany). Materials were obtained from the following manufacturers: 316L Stainless Steel (MetShap, Germany), KeraTi5 (Eisenbacher Dentalwaren, Germany), PM-AG101P (Legor Group, Italy), LithaLox 350 (Lithoz, Vienna), LithaCon ATZ 980 (Lithoz, Vienna), LithaCon 3Y 210 (Lithoz, Vienna), PEEK (Whitepeaks Dental Soulutions, Germany).

### Animals

Male 8–12 weeks old, 20–26 g, C57BL/6J-Tg(Thy1-GCaMP6s)GP4.12Dkim/J mice and C57BL/6J from Charles River Laboratories (Sulzfeld, Germany) were used. Mice were group-housed under pathogen-free conditions and bred in the animal housing facility of the Institute of Stroke and Dementia Research (Munich, Germany), with food and water provided ad libitum (21 ± 1 °C, at 12/12 h light/dark cycle). Animal husbandry, health screens, and hygiene management checks were performed in accordance with the Federation of European Laboratory Animal Science Associations (FELASA) guidelines and recommendations. All experiments were carried out in compliance with the National Guidelines for Animal Protection of Germany with the approval of the regional animal care committee of the Government of Upper Bavaria and were overseen by a veterinarian (Animal license numbers: Vet_02-21-37 and Vet_02_21_46).

### Cranial window implantation

For two-photon microscopy a chronic cranial window was implanted three weeks before intravital imaging. Mice were anesthetized by intraperitoneal injection of a medetomidine–midazolam–fentanyl (MMF) mixture (0.5 mg/kg medetomidine, 5 mg/kg midazolam, 0.05 mg/kg fentanyl). A circular craniotomy was performed over the right barrel cortex using stereotaxic coordinates relative to bregma (1.1 mm posterior, 3.3 mm lateral). The skull flap and dura were carefully removed to expose the cortical surface. A total volume of 0.350 µl of AAV9-Syn-jGCaMP8m-WPRE (Addgene viral prep #162375-AAV9) diluted at a 1:10 ratio in PBS to a final titer of 1 × 10^12^ vg/ml, was delivered into barrel cortex via a pulled glass capillary connected to a Nanoliter 2020 injector (World Precision Instruments) at 50 nl/min. The injection site was targeted at −1.1 mm rostrocaudal, −3.3 mm mediolateral, and −0.25 mm dorsoventral relative to bregma. Following virus injection, a 3 mm circular glass coverslip was placed over the craniotomy and sealed to the skull using histoacryl tissue adhesive. For subsequent head-fixed imaging, a custom zirconium head plate or a custom steel headplate (Femtonics Ltd., Budapest, Hungary) were mounted on the skull with UV-curable dental cement. At the end of the procedure, anesthesia was antagonized with an intraperitoneal injection of atipamezole (0.5 mg/ml), naloxone (3 mg/ml), and flumazenil (5 mg/ml). Animals were maintained overnight in a 32 °C heated chamber to support post-operative recovery and were allowed an additional two weeks of recuperation before *in vivo* imaging experiments were performed.

For mesoscale multimodal imaging a minimal-invasive cranial window was implanted one week before the first imaging session. Mice were anesthetized with isoflurane (5% induction, 2% maintenance) delivered in 70% N_2_O and 30% O_2_, positioned prone, and secured in a stereotaxic frame (51500, Stoelting Europe, Ireland). To prevent corneal drying, 5% dexpanthenol ointment (Bepanthen, Bayer, Germany) was applied to both eyes. After confirming surgical depth of anesthesia (absence of the foot-pinch reflex), the scalp was disinfected with a topical antiseptic (Octenisept, Schülke, Germany). The skin and overlying connective tissue were removed to expose the dorsal skull, and the periosteum was gently cleared. Local analgesic (1% Xylocaine) was applied to the exposed muscles and the wound edges. The skull surface was then cleaned with 60% ethanol and allowed to air-dry. A thin layer of transparent, biocompatible dental cement (Quick Base S398, L-Powder clear S399, Universal Catalyst S371, Parkell C&B Metabond, USA) was used to seat a curved coverslip (Crystal Skull, LabMaker, USA) and to secure a zirconium head-fixation frame around the window. The coverslip was carefully pressed into place to avoid trapping air bubbles and to ensure a minimal layer of cement. The head frame was held steady until initial polymerization (∼1 min), then the implant was allowed to cure for an additional 5–10 min. Following surgery, mice recovered in a warmed chamber for ∼30 min before being returned to their home cage and were monitored daily until imaging.

### Photothrombotic stroke model

For *in vivo* widefield calcium imaging, a photothrombotic (PT) stroke was induced in the left somatosensory cortex as described previously [27]. Rose Bengal (1% in saline; Sigma, 198250-5G) was administered intraperitoneally (10 µl/g body weight) 10 min before light exposure. Anesthesia was achieved with isoflurane (5% induction, 2% for surgical preparation, and 0.5% during illumination) in a carrier gas consisting of 30% O_2_ and 70% N_2_O. Once a stable surgical plane was reached, animals were positioned in a stereotaxic apparatus (51500, Stoelting Europe, Ireland), and core temperature was kept at 37°C using a feedback-controlled heating pad. To avoid corneal dehydration, dexpanthenol eye ointment (Bepanthen, Bayer, Germany) was applied to both eyes. Bregma was located through the transparent cranial window by identifying the coronal–sagittal suture junction, and the illumination site was defined in the left hemisphere at 1.5 mm lateral and 1.0 mm anterior to bregma. A removable cover was placed on the cranial window, leaving a circular opening (0.5 mm diameter) aligned to the target area. At 10 min after dye injection, the cortex was illuminated for 17 min using a 561 nm laser set to 25 mW output power (Cobolt Jive 50, Hübner Photonics, Sweden) coupled with an FC/APC fiber collimation unit (f = 7.66 mm) and a zero-aperture graduated iris (SM1D12CZ, Thorlabs, USA) adjusted to 0.5 mm. Following illumination, mice were returned to their home cage for recovery. Successful lesion induction was first assessed by visual detection of a focal cortical infarct through the cranial window.

### *In vivo* two-photon microscopy

*In vivo* two-photon imaging was conducted using a Leica SP8 DIVE multiphoton microscope equipped with a femtosecond pulsed laser, a 25×/1.0 NA (Leica Microsystems, Germany) water-immersion objective, and a motorized stage. GCaMP8m fluorescence was excited at 940 nm, and emitted photons were collected between 500 and 550 nm using non-descanned detectors positioned in the epifluorescence detection path. The laser power at the sample was maintained at approximately 50 mW to limit photodamage. Time-lapse image series of GCaMP8m-expressing neurons were acquired at 27.84 Hz with a frame size of 512 × 512 pixels and an axial step of 1 µm at imaging depths of 200–250 µm below the pial surface. After an initial baseline recording of approximately 3 minutes, a 20 s, 5 Hz stimulation and 80 ms pulse duration paradigm was delivered via Picospritzer (Parker-Hannifin Corporation) triggered via an Arduino-based control system. This stimulation protocol was delivered four times per session, with successive stimulation epochs separated by ∼3 minutes of spontaneous activity.

### *In vivo* widefield multimodal imaging and sensory stimulation

To monitor calcium activity in layer 2/3 cortical excitatory neurons C57BL/6J-Tg(Thy1-GCaMP6s)GP4.12Dkim/J [28] transgenic calcium indicator mouse line was used. Before awake imaging, mice completed 5 consecutive days of habituation to the gramophone head-holder (Gramophone, Femtonics Ltd., Hungary) and handling procedures. On imaging days, animals were briefly anesthetized with isoflurane (4%) to attach the pre-implanted head holder to the headpost and were then allowed to fully recover before data acquisition. Head-fixed mice were imaged in a darkened environment that permitted unrestricted body movement while maintaining stable cranial fixation. Movement of the Gramophone platform was recorded directly from the head-fixation platform as acceleration (±) using a Python script provided by Femtonics. Infrared behavioral video was acquired in parallel using a monochrome low-light camera (ace2A1920, Basler AG, Germany) equipped with a 2/3″ C-mount lens (C23-1216-2M, Basler AG, Germany). A 3D-printed eye shield was used to prevent stray illumination from reaching the eyes. Mice underwent an initial pre-acquisition period (2 min) followed by a 4-min imaging acquisition. Fluorescence and intrinsic optical signal (IOS) reflectance were collected with a high-speed sCMOS camera (Andor Zyla 5.5, Oxford Instruments, UK) coupled to a video objective (AF D Mikro 60/2.8, Nikon, Japan). Illumination was provided by a custom triple-LED module delivering 470 nm, 530 nm, and 625 nm light (M470L4, M530L4, M625L4; Thorlabs, USA). Each LED was equipped with the corresponding laser-line filter (FL488-10, FLH532-4, FLH635-5; Thorlabs, USA), collimation optics (ACL2520U-A, SM1T-2; Thorlabs), and modular optomechanics (Thorlabs, USA). Emission light was collected through a 515 nm long-pass filter (FGL515M, Thorlabs, USA) such that 470 nm excitation generated the fluorescence channel (blue), while 530 nm (green) and 625 nm (red) illumination yielded two IOS reflectance channels. Illumination was synchronized to the camera TTL trigger and controlled by an Arduino microcontroller (UNO, Arduino, Italy) programmed to strobe LEDs sequentially on each frame (no illumination, blue, green, red). The system covered a 10 × 10 mm field of view sampled at 512 × 512 pixels. Recordings were acquired at 100 Hz for 4 min, generating 16-bit raw image stacks comprising fluorescence, two IOS reflectance channels, and non-illuminated background frames. All data were transferred to the analysis workstation for offline processing.

### Microscope image data analysis

Time-lapse two-photon calcium imaging movies were processed using established software for denoising, motion correction, and region-of-interest (ROI) extraction. Raw fluorescence videos were first denoised with DeepCAD-RT [29]. The denoised movies were then processed with Suite2p [30] to perform motion correction, quantify framewise x–y displacements as a measure of motion, and correct for motion artifacts. Suite2p was further used to segment active neuronal regions of interest and to extract fluorescence time courses for each ROI for subsequent analyses. For each neuron, a baseline fluorescence value (F_0_) was calculated as the mean fluorescence in a 2-second window prior to stimulus onset. The change in fluorescence relative to baseline (ΔF/F_0_) was then calculated as (F – F_0_) / F_0_. Extended peri-stimulus fluorescence traces spanning 10 s before and 20 s after stimulus onset were extracted for each stimulation epoch. Responses were analyzed independently for each neuron and each of the four stimulation trials, such that a single neuron could contribute multiple responsive trials. For every trial, baseline fluorescence (F0) was calculated from the 2 s period preceding stimulus onset, and relative fluorescence changes were computed as ΔF/F0 = (F – F0)/F0. Neurons were classified as responsive when the peak ΔF/F0 during stimulation exceeded four standard deviations above baseline activity. All responsive neuron–stimulation trials were pooled for population-level analyses. To quantify stimulus-evoked activity, mean ΔF/F0 values were calculated for two temporal windows in each responsive trial: a 2 s baseline period and a 20 s window beginning at stimulus onset. Mean fluorescence values from these windows were extracted for each trial and used for paired statistical comparisons. Differences between baseline and post-stimulus activity were assessed using paired two-tailed Student’s *t*-tests implemented in SciPy (Python).

Widefield multimodal imaging data were processed as previously described workflows [26, 31] with additional custom analyses. Image preprocessing began with channel separation, followed by rigid motion correction by registering each frame to a whole-recording average reference image. Transformations were estimated from the non-illuminated background channel using *imregtform* algorithm, and the resulting frame-wise transformation matrices were applied to all channels (fluorescence and reflectance) using *imwarp* algorithm. Frame-wise geometric transformations were then converted from pixels to physical units using the imaging scale (35.5 µm/pixel), yielding residual in-plane drift estimates in the mediolateral (x) and anteroposterior (y) directions for each frame. Drift parameters were summarized per recording and compared across experimental sessions, together with the Gramophone-derived motion readout, expressed as velocity computed from the absolute acceleration signal (i.e., magnitude-only, independent of movement direction; Fig. S3B and Movie S2). Recordings were then manually aligned across sessions (within and between recording days) using translation-only registration implemented in a custom MATLAB GUI. Rolling background subtraction was performed by subtracting, for each fluorescence and reflectance frame, its corresponding non-illuminated background frame. ΔF/F was computed on a per-pixel basis by first centering the time series and then normalizing it to the pixel-wise mean fluorescence over the full recording. To correct hemodynamic contamination, oxy- and deoxyhemoglobin traces (ΔHbO, ΔHbR) were derived from the two reflectance channels and used for hemodynamic correction in custom C# code (.NET 8.0). In practice, applying both excitation- and fluorescence-based corrections caused hemodynamic signals to dominate the corrected fluorescence; therefore, we omitted excitation correction and assumed constant fluorophore illumination across trials. Corrected fluorescence signals were then low pass filtered (Chebyshev, 5 Hz cutoff) and down sampled to 10 Hz. To mitigate filter edge artifacts, the first and last 10 s of each recording were discarded. All preprocessing steps were performed in MATLAB (MathWorks R2016b, 2025a).

Functional connectivity was quantified by computing Pearson correlation coefficients between signal time courses extracted from 16 anatomically and functionally distinct cortical ROIs (bilateral primary motor, somatosensory, and visual–associational regions). ROIs were defined using independent vector analysis on baseline resting-state recordings [26] and cross-referenced with a data-driven iterative parcellation approach [32]. Correlation coefficients were Fisher z-transformed for group comparisons. Group differences were visualized as topographic maps across the dorsal cortical surface. For multimodal visualization, functional maps were rigidly registered and scaled to a 3D cortical surface reconstruction derived from T2-weighted MRI (Imaris 10.2, BitPlane, Oxford Instruments, UK), using the brain surface boundaries as anatomical constraints.

### MRI imaging

Three Tesla MRI was performed on a small-animal scanner (3T nanoScan PET/MR, Mediso, Hungary) one day before and one day after sham or photothrombotic stroke induction. For scanning, mice were anesthetized with isoflurane (4% induction, 1.2% maintenance) delivered via face mask in a 30% O_2_ / 70% N_2_O gas mixture. Respiratory rate and body temperature (maintained at 37 ± 0.5 °C) were continuously monitored using an abdominal pressure-sensitive pad and a rectal temperature probe, and anesthesia levels were adjusted to keep physiological parameters within the normal range.

The following sequences were acquired: (I) coronal T1-weighted imaging using a 2D gradient echo (GRE) sequence (TR/TE = 200/3.2 ms; flip angle = 35°; FOV = 50 × 45 mm; in-plane resolution = 250 × 250 µm; 36 slices; slice thickness = 400 µm), and (II) axial T2-weighted imaging using a 3D spin-echo sequence (TR/TE = 2500/91.3 ms; FOV = 51 × 52 mm; in-plane resolution = 203 × 203 µm; 214 slices; slice thickness = 200 µm). MRI data were post-processed in ImageJ and subsequently aligned to the widefield-derived functional connectivity maps in MATLAB using translation and isotropic scaling; overlays were generated in MATLAB.

11.7 Tesla MRI measurements were conducted using a Bruker BioSpec 117/16 USR system operating at 11.7 T (16 cm free bore), equipped with a B-GA9S gradient system. Signal reception was performed using a 1H MRI CryoProbe two-element phased-array coil designed for mouse imaging (Bruker BioSpin MRI GmbH, Ettlingen, Germany). To evaluate the MRI compatibility of the investigated materials, samples were embedded in 2% (w/v) low-melting-point agarose (AppliChem, Germany) within 15 ml falcon tubes (Greiner Bio One, Germany). All 3D printed samples were manufactured with identical geometry and dimensions. Three-dimensional datasets of the embedded samples were acquired using a T1-weighted isotropic Fast Low-Angle Shot (FLASH) pulse sequence (field of view [FOV] = 15 × 15 × 15 mm^3^; acquisition matrix = 256 × 256 × 256, yielding an isotropic in-plane resolution of 58 μm; repetition time [TR] = 50 ms; echo time [TE] = 6.15 ms; flip angle = 15°; number of signal averages = 3). The acquired MRI data were converted to DICOM format for post-processing in Fiji. Subsequently, 3-dimensional data sets were color inverted, filtered using a Sobel edge detector and projected in z-direction.

### Micro–computed tomography (micro-CT)

Samples were imaged using a high-resolution micro–computed tomography system (SkyScan 1273, Bruker microCT, Kontich, Belgium). Scans were performed using a Hamamatsu L9181-02 X-ray source and a Varex 2315 flat-panel detector, operated at a source voltage of 130 kV and a current of 115 µA. Projection images were acquired with an isotropic voxel size of 8.0 µm, using a camera binning of 1 × 1 and an exposure time of 554 ms per projection.

For each scan, 2401 projection images were collected over a full 360° rotation with a rotation step of 0.15°. Frame averaging (6 frames per projection) and flat-field correction were enabled to improve signal-to-noise ratio. Scans were performed using a step-and-shoot acquisition mode with counterclockwise gantry rotation. The object-to-source distance was 53.6 mm and the camera-to-source distance was 501.0 mm.

Raw projection images were reconstructed using NRecon software (version 1.7.5.1, Bruker microCT) with GPU-based reconstruction. Reconstruction was performed over a 360° angular range using a Hamming filter (α = 0.54) with a cutoff at 100% of the Nyquist frequency. Reconstructions were generated as 16-bit grayscale images with an isotropic voxel size of 8.0 µm. A region of interest (ROI) was defined during reconstruction to include the complete specimen while excluding empty field-of-view regions.

### Quantification and Statistical analysis

Data were analyzed using GraphPad Prism version 9.0 and 10. All summary data are expressed as the mean ± standard deviation (s.d.), unless indicated otherwise. Normality was assessed in all datasets using the Shapiro-Wilk normality test. Normally-distributed data were analyzed using a two-way Student’s t test (for 2 groups) or ANOVA (for > 2 groups). Non-normally distributed data were analyzed using the Mann-Whitney U test (unpaired data) or Wilcoxon rank sum test (paired data) (for 2 groups), or Kruskal-Wallis test (H test, for > 2 groups). Multiple comparison adjusted p values were computed using Bonferroni correction or Dunn’s multiple comparison tests. For all analyses a p value < 0.05 was considered statistically significant.

## Acknowledgements

The authors thank Prof. Dr. Jürgen Bernhagen for providing technical support with the Leica SP8 DIVE. S.F was funded through the “iBehave” research consortium (https://ibehave.nrw/) from the program “Netzwerke 2021” an initiative of the Ministry of Culture and Science of the State of North Rheine-Westphalia, Germany. F.C.N. received funding from the German Research Foundation DFG (SPP2395) and the CANTAR network (funded by the Ministry of Culture and Science of the State of North Rhine-Westphalia; the funders had no role in study design, data collection, and interpretation, or the decision to submit the work for publication). GP-AAV-syn-jGCaMP8m-WPRE was a gift from GENIE Project (Addgene plasmid #162375; http://n2t.net/addgene:162375 ; RRID:Addgene_162375). Fig. 2A was created with Biorender.

## Authors contribution

S.F., G.M.C., D.V., H.D., and C.H. designed and performed the experiments, and acquired, processed, and analyzed the data. S.F., G.C., D.P.V., and H.F. drafted the manuscript. H.F. conceptualized the study, designed and supervised the experiments, and analyzed and interpreted the data. P.M. designed the experiments and analyzed the data. B.E.-D. acquired, processed, and analyzed the data. F.N., W.W., A.L., P.S., D.H., and E.F. conceptualized the study. All authors critically revised the manuscript.

## Data and Material Availability

All data are available in the main text or the supplementary materials. Custom written scripts for bioinformatic analysis and headholder CAD file will be made available in a GitHub repository upon publication. Any additional information required to reanalyze the data reported in this paper is available from the lead contact upon request.

