## Supplemental Figures for "MRI-Compatible Rigid Head Holders for Artifact-Free Multimodal Imaging in Mice"

### Supplementary material list:

- Supplementary Figures S1-S4

Figure S1: Sketch of 3D printed zirconia head holders.
Figure S2: Two-photon imaging confirms whisker-evoked calcium responses in cortical neurons.
Figure S3: Quantification of image motion and locomotor activity during mesoscale imaging.
Figure S4: Atlas-referenced coronal MRI view of stroke-associated edema 1 day after photothrombotic stroke.

- Supplementary Table 1: Overview of the elastic modulus and density values of tested materials.
- Supplementary Movie S1: Representative two-photon calcium imaging from the somatosensory cortex in a naive awake, head-fixed mouse.
- Supplementary Movie S2: Representative multimodal mesoscale imaging in a naive awake, head-fixed mouse.
- Supplementary Movie S3: Three-dimensional MRI reconstruction with Imaris surface segmentation 1 day after photothrombotic stroke.

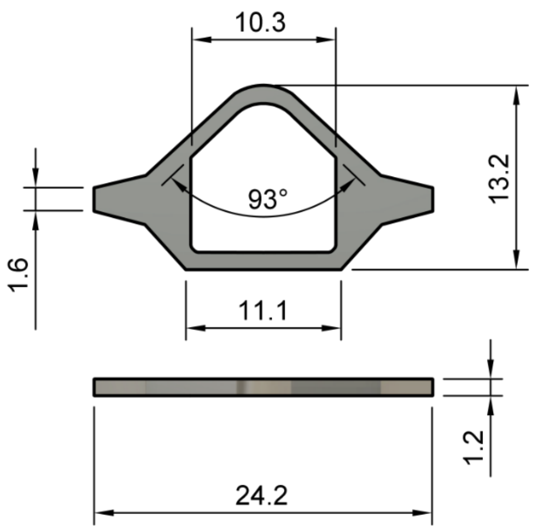


**Suppl. Figure 1:** Construction sketch of 3D printed zirconia head holders used for *in vivo* 2-photon, mesoscale and MRI imaging. Numbers are given in mm.

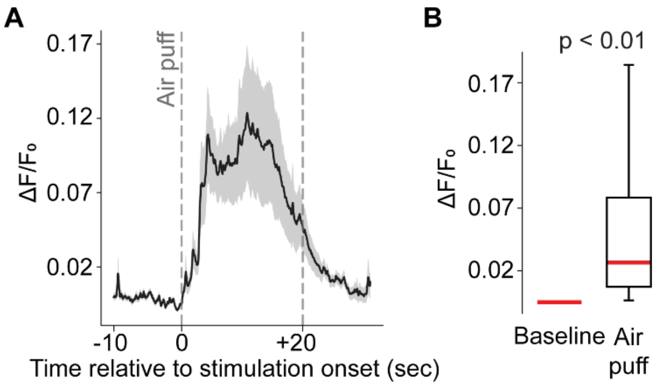


**Suppl. Figure 2: (A)** Normalized population calcium responses across the four whisker stimulation trials. Data are shown as mean ± SEM (n = 654 responses from 236 cells). **(B)** Quantification of mean ΔF/F0 calculated during baseline and stimulation period (four stimulation trials). Data are presented as median and interquartile range (25th–75th percentile). Whiskers indicate the full range of the data (minimum to maximum). Statistical comparison was performed using a paired Student's *t*-test.

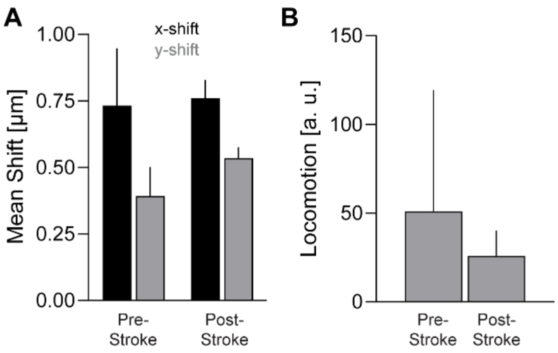


**Suppl. Figure 3:** Quantification of image pixel shift and locomotor activity during mesoscale imaging before and after photothrombotic stroke. **(A)** Bar plots show quantitative measurements of x- and y-axis motion during image acquisition before and after stroke induction. Bars represent mean ± SD of the average pixel shift converted to micrometers across the full recording period. **(B)** Bar plots show mean ± SD locomotor activity recorded from the rotating disc treadmill during image acquisition before and after stroke induction.

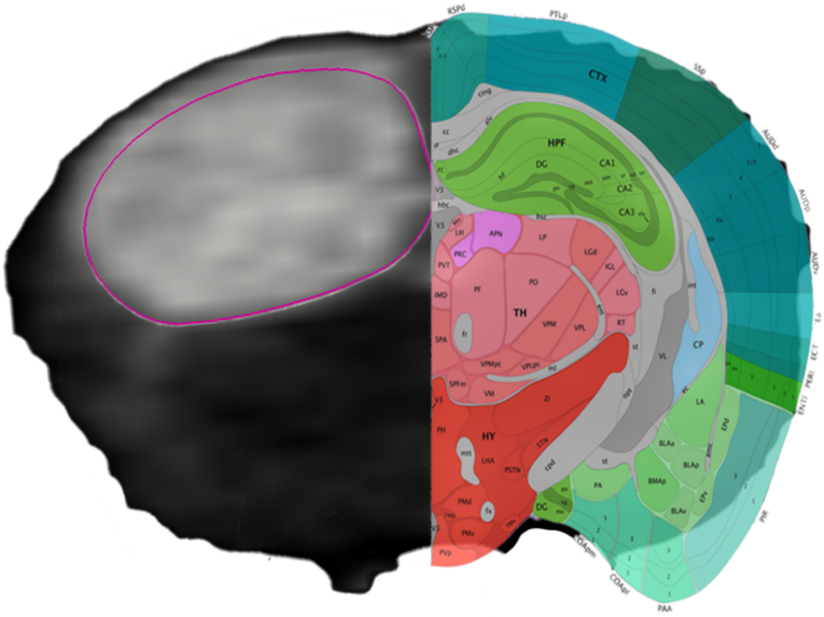


**Suppl. Figure 4:** Atlas-referenced coronal MRI view of stroke-associated edema 1 day after photothrombotic stroke. T2-weighted MRI acquired 1 day after photothrombotic stroke shows pronounced hyperintensity in the left hemisphere, consistent with edema. The matching coronal plane from the Allen Mouse Brain Atlas, Mouse P56 coronal reference atlas image 77, is overlaid on the contralateral hemisphere to provide anatomical reference. The edema extends beyond the cortical induction site and includes hippocampal, dorsal *caudoputamen*, and thalamocortical regions.

| | Elastic Modulus (GPa) | Density ($g/cm^3$) |
|---|---|---|
| **Steel** | 176 | n.d. |
| **Titanium** | 112 | 4.4 |
| **Silver** | n.d. | 10.4 |
| **Alumina** | 380 | 3.9 |
| **AtZ** | 220 | 5.5 |
| **Zirconia** | 205 - 210 | 6 |
| **PEEK*** | 4 | 1.3 |

*Limaye *et al.*, Heliyon, 2022

**Suppl. Table 1:** Overview of the elastic modulus and density values of tested materials.
